# Anti-S-layer monoclonal antibodies impact *C. difficile* physiology

**DOI:** 10.1101/2023.09.21.558785

**Authors:** Lise Hunault, Emile Auria, Patrick England, Julien Deschamps, Romain Briandet, Vanessa Kremer, Bruno Iannascoli, Léo Vidal-Maison, Chunguang Guo, Lynn Macdonald, Séverine Péchiné, Cécile Denève-Larrazet, Bruno Dupuy, Guy Gorochov, Pierre Bruhns, Delphine Sterlin

## Abstract

*Clostridioides difficile* (*C. difficile*), a gram-positive anaerobic and spore-forming bacterium, is the leading cause of nosocomial antibiotic-associated diarrhea in adults and is characterized by high levels of recurrence and mortality. Surface-layer Protein A (SlpA), the most expressed protein on bacterial surface, plays a crucial role in the early stages of infection although its role in *C. difficile* physiology is yet to be fully understood. Anti-S-layer antibodies have been identified in the sera of convalescent patients and correlate with improved outcome of *C. difficile* infection (CDI). However, the precise mechanisms of how anti-S-layer antibodies can confer protection to the host remain unknown. In this study, we report the first monoclonal antibodies (mAbs) targeting S-layer of the reference strain 630. Characterization of these mAbs unravels important roles for S-layer protein in growth, toxin secretion, and biofilm formation with, surprisingly, opposite effects of different anti-SlpA mAbs on these functions. One anti-SlpA mAb impaired *C. difficile* growth and restored sensitivity to lysozyme-induced lysis. These findings suggest that anti-S-layer antibody responses may include protective and detrimental effects for the host and provide important insights for designing adequate S-layer-targeting therapeutics.

## Introduction

*Clostridioides difficile* is an anaerobic, gram-positive, spore-forming bacteria, that is the leading agent responsible for nosocomial antibiotic-associated diarrhea and colitis in adults^1^. *C. difficile* infection (CDI) causes substantial morbidity and mortality with severe pseudomembranous colitis characterized by extensive colonic damage and intestinal inflammation. While CDI symptoms have largely been attributed to the bacterial toxins, a growing concern focused on *C. difficile* adhesins and surface proteins involved in gut colonization and evasion of the immune system surveillance. These proteins play a major role in triggering bacterial pathogenesis through interactions with Toll-Like Receptor 4 (TLR4) and inflammatory response induction^1,2^. Among these proteins, *C. difficile* Surface-layer protein A (SlpA) has gained substantial interest.

The *C. difficile* S-layer is composed of two main proteins *i.e.,* the High-Molecular Weight (HMW) and the Low-Molecular Weight (LMW) Surface Layer Proteins (SLPs) that derive from the common precursor SlpA. SlpA is first secreted and then cleaved by the cell wall cysteine protease Cwp84, releasing the two mature subunits HMW and LMW. These two subunits associate to form a stable heterodimeric complex, which is anchored to the cell wall by the HMW, with the LMW being the most external subunit. SlpA is secreted throughout the cytoplasmic membrane and constitutes an interwall reservoir, which is available to fill the gaps that form during growth or damage^3^. With the assembly of the S-layer at areas of newly synthesized peptidoglycan, *C. difficile* can maintain a stable S-layer that continually protects the cell. One astonishing characteristic of *C. difficile* S-layer is its compactness. With pores of only 10Å in diameter, it is more compact than other S-layers whose pores range from 30Å up to 100Å. This renders *C. difficile* impermeable to large molecules such as lysozyme^4^, to which it is resistant.

The S-layer is crucial for bacterial integrity, and *C. difficile* S-layer-null mutants display severe impaired physiological functions. They are highly sensitive to innate immune effectors such as lysozyme, show sporulation defects, and produce less toxins *in vitro*^5^. *C. difficile*’s persistence and recurrence were linked to the presence of spores^6^ and suggested to be associated to its ability to form biofilms in the gut^7^. Biofilm formation is the differential process of planktonic cells to bacterial communities embedded into a thick enclosed matrix^8^. Cwp84 mutants with altered S-layer display an increased biofilm generation suggesting that intact S-layers prevent aggregation, which is one of the first steps to generate biofilms^9^. As the predominant surface protein, *C. difficile* S-layer has also been implicated in attachment to intestinal cells both *in vitro* and *ex vivo*^10,11^.

The S-layer is immunogenic, as anti-SLPs antibodies have been detected in the sera of convalescent patients and are associated with improved CDI outcome^12,13^. In animal models, passive immunization using anti-SlpA serum has been demonstrated to delay *C. difficile* colonization in mice^14^, whereas active immunization with recombinant SlpA slightly prolonged survival of hamsters infected by *C. difficile*^15^. Additionally, anti-LMW nanobodies have been shown to decrease bacterial motility *in vitro*^16^. However, the extent to which anti-S-layer humoral responses interfere with *C. difficile* fitness and CDI pathogenesis remains unclear. No monoclonal antibodies (mAbs) targeting the S-layer that could be used to explore the role of SlpA *in vivo* have been reported so far.

Here, we generated and characterized the first anti-LMW mAbs to interrogate S-layer interactions with host immune response. We describe differential effects of anti-LMW mAbs on *C. difficile* physiology in terms of growth, toxin secretion, and biofilm formation *in vitro*. Our work deciphers interactions between antibodies and various epitopes of the S-layer with unexpectedly different outcomes and describes further the role of *C. difficile* S-layer in bacterial fitness.

## Results

### Generation and characterization of high-affinity LMW-specific mAbs

To interrogate the role of the S layer in *C. difficile* biology, we generated a collection of mAbs targeting the SlpA LMW of the reference *C. difficile* 630Δerm (CD630Δ*erm*)-a spontaneous erythromycin sensitive derivative of the reference strain 630, the most external subunit of the S-layer. As anti-LMW antibodies may potentially be of therapeutic interest for the treatment of *C. difficile* infections, we used knock-in mice in which the endogenous genes encoding the heavy chain variable domain (VH) and the kappa light chain variable domain (Vk) were replaced by their human counterparts (Velocimmune)^17,18^ with one modification: only one allele of the endogenous Vk locus was replaced by human Vk segments, and the second allele of the endogenous Vk locus was replaced by human Vl segments (Supplemental Fig. 1a). As the Vk locus expresses 95% of the light chains in mice ^19^, placing human Vl segments at the Vk locus increases the variability of light chain expression. Thus, after hybridoma identification, cloning of these VH and VL into vectors containing human heavy and light chain constant domains, allows for direct development - *in fine* – of fully human anti-LMW mAbs. These mice but also BALB/c mice were immunized with recombinant LMW at D0, D21, D42 and four days before spleen collection, according to the schedule presented in Fig. 1a. Anti-LMW hybridomas were generated from splenocytes of one Velocimmune and one BALB/c mouse, using ELISA as a screening method (Fig. 1a). Seven anti-LMW mAbs (all mouse IgG1) were identified that demonstrated a 10^-1^ to 10^-2^ µg/mL effective concentration 50 (EC_50_) in an anti-LMW ELISA. mAbs NF10 and KH2 originated from the BALB/c mouse and possess mouse VH-VL sequences, whereas mAbs 1E2, 2B7, 2C4 and 4G4 originated from the Velocimmune mouse and possess human VH-VL sequences. For all mAbs, their VH-VL gene sequences displayed CDR3 length distributions from 10 to 20 residues (Table S1).

**Figure 1:**
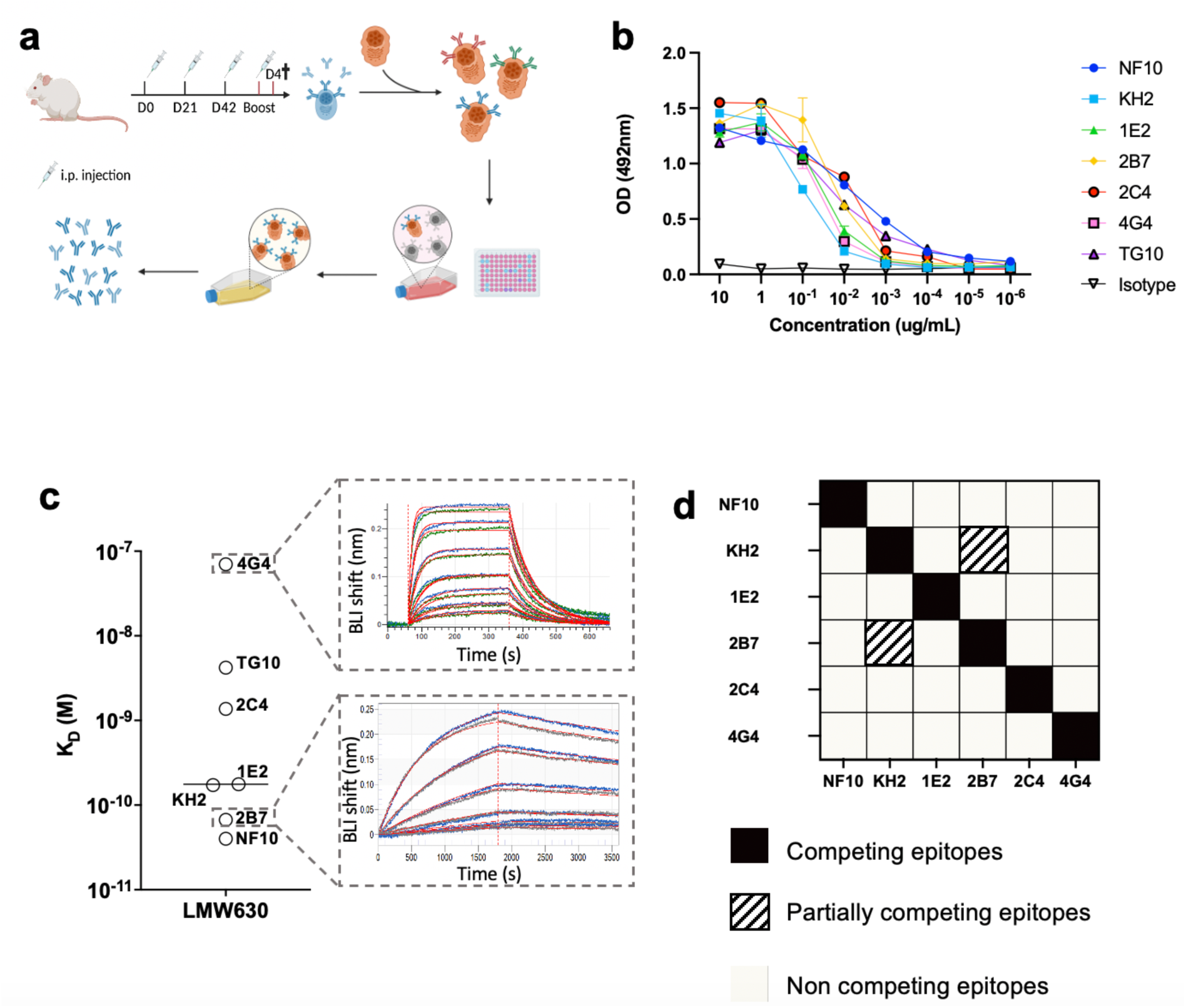
High-affinity anti-LMW mAbs bind distinct epitopes. **a.** Schematic view of immunization, hybridoma generation and screening for obtention of anti-LMW mAbs. **b.** Mab binding to recombinant LMW measured by ELISA at indicated concentrations. Dark curve represents isotype control. **c.** Affinities towards LMW determined by Bio-Layer Interferometry. Representative sensorgrams of one low (4G4) and one high-affinity (2B7) mAb. Antibody concentration from 500 nM to 8 nM for 4G4 and from 2 nM to 0.02 nM for 2B7 were tested, as shown from top to bottom. Blue curves represent raw data while red curves represent fitting with a 1:1 antibody:antigen model. **d.** Summary table representing the results of BLI-based competitive of anti-LMW mAbs towards LMW.

Bio-layer interferometry (BLI) experiments revealed a very large range of equilibrium dissociation constants (K_D_) ranging from 32 pM to 70 nM, corresponding to low to very-high affinity antibodies (Fig. 1c). The mAb with the worse affinity displayed a fast on/off profile with a high dissociation rate (k_off_) of ∼0.01 s^-1^, whereas the two mAbs with the best affinities displayed a very low k_off_ of ∼0.00003 s^-1^ (Table S1). To examine whether anti-LMW mAbs recognized overlapping or distinct epitopes on LMW, we designed a competitive BLI assay based on a pre-bound anti-LMW Ab as a competitor. Only two mAbs, KH2 and 2B7, partially competed for their binding to LMW (Fig. 1d). We therefore generated a set of mostly high-affinity anti-LMW mAbs that target 5 different and non-overlapping epitopes on *C. difficile* SlpA LMW-630.

### Binding to *C. difficile* 630 vegetative cells

Since the LMW is the most exposed S-layer protein of *C. difficile*, we next wanted to assess mAb binding to *C. difficile* whole bacteria. For this purpose, we used a previously reported bacterial flow cytometry assay^20^. Five out of the seven anti-LMW mAbs readily bound CD630Δ*erm* (Median Fluorescence Intensities (MFI) 100- to 1,000-fold higher compared to isotype control). mAb 2C4 poorly bound *C. difficile* strain 630 (MFI 5-fold higher compared to isotype) and mAb 4G4 very poorly if not at all (MFI 2.5-fold higher compared to isotype) (Fig. 2a). These results are mostly in agreement with the affinities of these mAbs for LMW, as mAb 4G4 possesses by far the worst affinity (70nM). mAb 2C4, however, should bind *C. difficile* in these conditions (K_D_= 1.37nM) but its epitope may be partially inaccessible. Also, 2B7 that possesses a very high affinity (K_D_= 67pM) displayed only a mild binding, 10x lower than that of NF10 that displays a similar affinity for LMW (K_D_= 43pM). None of these 7 mAbs cross-reacted with commensal bacteria of the same genus, i.e. *Clostridium bifermentans* and *Clostridium butyricum*, confirming their *C. difficile* specificity. In addition, none cross-reacted with a different ribotype (012) of *C. difficile* strain CD20-247, consistent with the low inter-strain homology of the LMWs (Fig. 2a).

**Figure 2:**
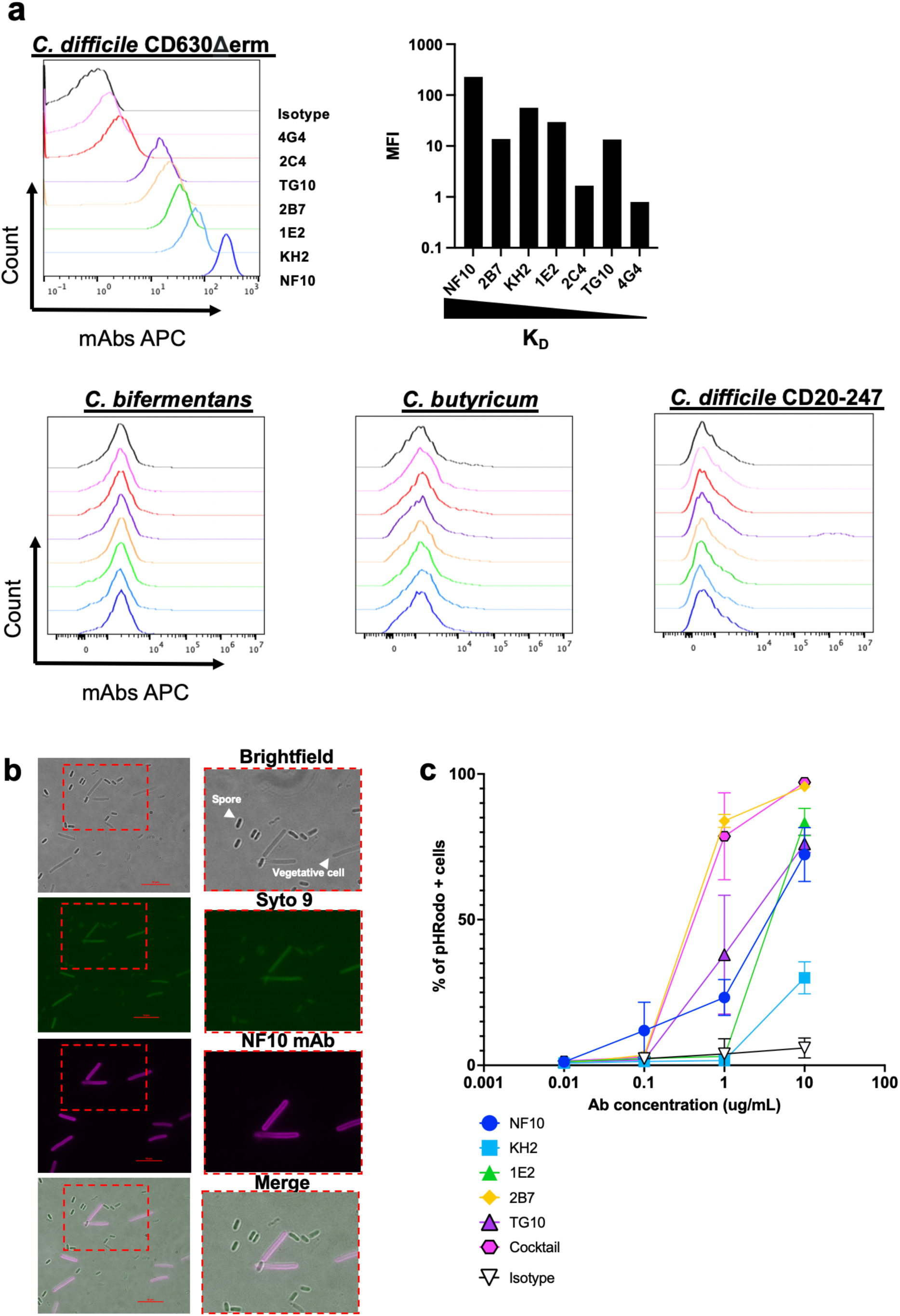
Anti-LMW mAbs bind vegetative *C. difficile* cells and enhance phagocytosis. **a.** Flow cytometry analysis of mAbs binding to indicated *C. difficile* strains and other *Clostridium* species (CD20-247 R012). Black curve corresponds to isotype control. **b.** Representative view of mAb binding to *C. difficile* vegetative cells but not to spores. DNA from vegetative cells and spores was labeled with SYTO9 while mAb-coated bacteria were stained with AF647-conjugated anti-mouse IgG antibody. Merged staining was presented on the right panel. Analysis was performed by confocal microscopy. **c.** Percentage of neutrophils that have phagocytosed *C. difficile-*opsonized by the indicated mAb or a cocktail of mAbs NF10, KH2, 1E2, 2B7 and TG10 at equimolar ratio, after 60 min and assessed by flow cytometry. Data represent mean + SEM of n = 3 technical replicates. Experiment was performed with at least 2 biological replicates.

### LMW is expressed at the surface of vegetative forms, but not spores

SlpA is expressed in the proteome of *C. difficile* spores, but whether the protein is exposed at the spores’ surface remains unknown^21^. We therefore analyzed by microscopy the binding of the mAb with the best K_D_ and the highest MFI on bacteria i.e., mAb NF10, to spores as well as to the vegetative form of *C. difficile*. Anti-LMW mAb NF10 stained the vegetative form but did not stain spores (Fig. 2b), suggesting that SlpA LMW is not similarly exposed on the surface of *C. difficile* spores.

### Anti-LMW mAbs enable *C. difficile* phagocytosis by neutrophils

We next evaluated if SlpA LMW was a suitable target for enabling or increasing phagocytosis of *C. difficile* by neutrophils, as it might occur during CDI after epithelial breakdown by the toxins secreted by *C. difficile*^22^ and invasion of the intestinal vili by bacteria and neutrophils^23^. We used a standard *in vitro* phagocytosis assay measured by flow cytometry in which bacteria are fluorescently labeled, opsonized by anti-bacterial IgG mAbs and incubated with purified human neutrophils. All anti-LMW mAbs being of the mouse IgG1 isotype, they are able to interact with human IgG receptors (FcγRs)^24^ expressed by human neutrophils. As expected, we found that binding of all five anti-LMW mAbs with a significant MFI on bacteria (excluding mAbs 4G4 and 2C4 from this analysis) enabled neutrophil-dependent phagocytosis of *C. difficile* (Fig. 2c). Surprisingly, we found no correlation between phagocytosis and staining by flow cytometry with mAb KH2, which exhibited a strong binding to *C. difficile* but resulted in minimal phagocytosis.. mAb 2B7 induced as much phagocytosis than a cocktail of mAbs NF10, KH2, 1E2, 2B7 and TG10 at equimolar ratio, suggesting a unique property of 2B7 or of its epitope to favor phagocytosis. Altogether, these results demonstrate that this set of five anti-LMW mAbs recognized *C. difficile* in a vegetative state and enhanced its phagocytosis by neutrophils.

### *C. difficile* growth is inhibited solely by mAb NF10

The S-layer appears to be essential for *C. difficile* fitness as *de novo* S-layer proteins should be assembled during cell growth and division^3^. We investigated if targeting of SlpA LMW may impact bacterial growth. When growth was measured in suspension, mAb NF10 strongly impacts *C. difficile* growth that only reach ∼50% of the plateau at 13 hours of culture when compared to that of the isotype control (Fig. 3a). No other anti-LMW mAb had an effect on growth. A minimum concentration of 50 µg/mL mAb NF10 was necessary to detect a statistically significant effect on growth (Supplemental Fig. 1b). The effect of NF10 mAb was specific to the *C. difficile* strain CD630Δ*erm* as no effect was detected with a *C. difficile* UK1 strain belonging to ribotype 027 (Fig. 3b). These results underline a unique property of mAb NF10 or of its epitope to inhibit growth of *C. difficile* strain CD630Δ*erm*.

**Figure 3:**
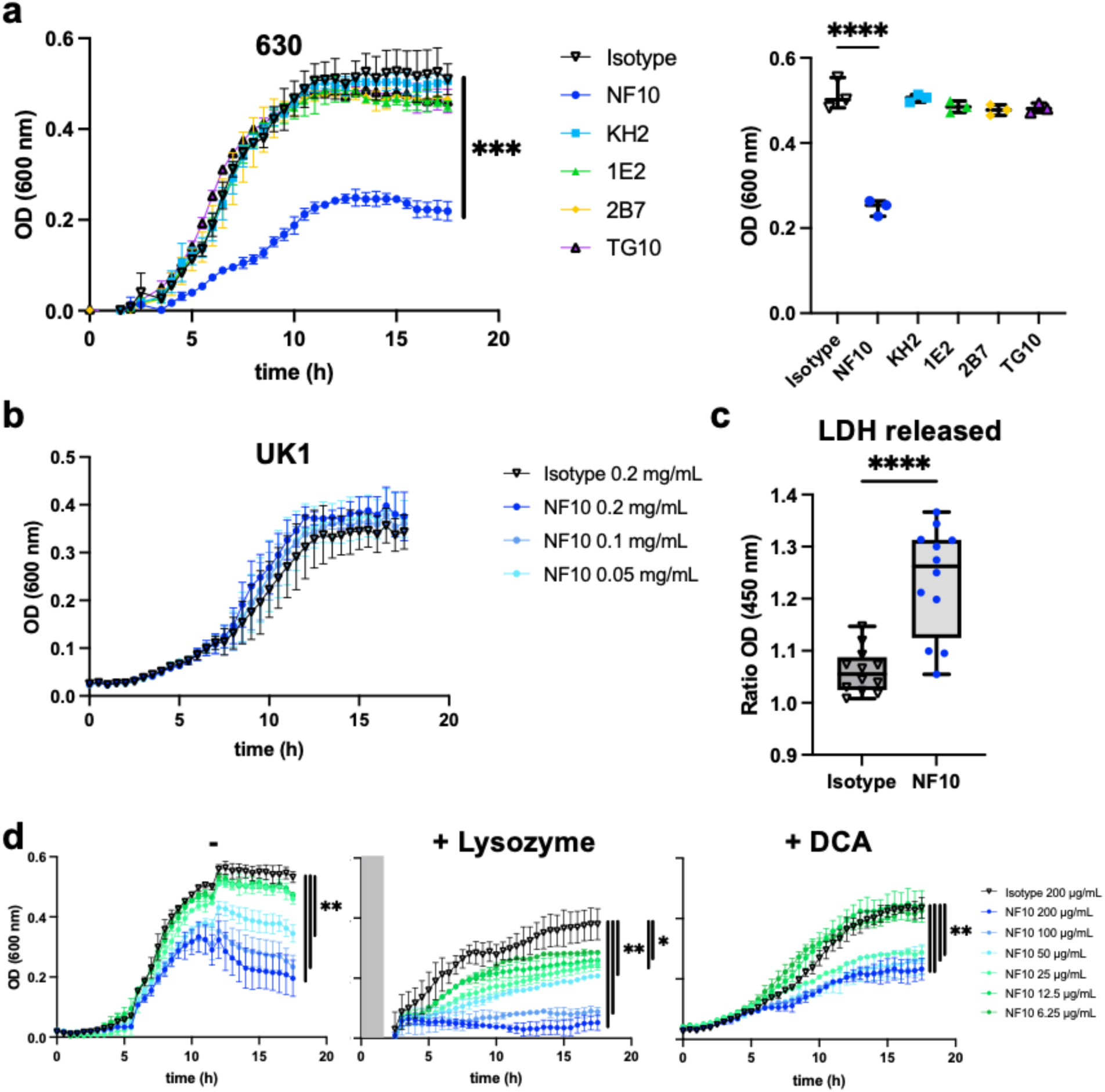
Effect on growth of anti-LMW mAbs and sensitivity to lysozyme and DCA. Cultures of *C. difficile* 630Δerm were inoculated at an OD_600nm_ of 0.05 and grown anaerobically at 37°C with OD_600nm_ measurements every 30 min. **a.** Effect of anti-LMW mAbs was assessed on growth. Left panel represents growth curves until 18h with measurements every 30 min for all anti-LMW mAbs and isotype. Right panel represents quantitative analysis at 13h for all anti-LMW mAbs and isotype. **b.** Effect of NF10 mAb was assessed on *C. difficile* UK1 strain growth at different concentrations. Data are presented as means and standard deviations from three technical replicates. **c.** LDH activity in the supernatant was normalized to condition without antibodies. The interquartile boxplots show medians (middle line), and the whiskers indicate minimal and maximal values. Asterisks indicate statistical significance calculated with a one-way ANOVA test followed by a Dunnett’s multiple comparison test (****p < 0.0001). Experiments were performed with two biological replicates in six technical replicates. **d.** Cultures of *C. difficile* 630Δerm incubated with different concentrations of NF10 mAb were monitored in combination with lysozyme (500 μg/ml), which was added after 2.5h growth or DCA (240 µM). Isotype control (dark lines) was included in all experiments. Data are presented as mean values (±SD) from three technical replicates. Asterisks indicate statistical significance with a two-way ANOVA test (ns: not significant; * p < 0.05, ** p < 0.01, *** p < 0.001, and **** p < 0.0001).

### Bacterial lysis is promoted by mAb NF10

We next sought to determine how anti-LMW mAb NF10 impaired *C. difficile* growth. A pool of SlpA precursor was reported to be localized within the bacterial cell wall, available to repair openings in the S-layer during cell growth or damage^25^. We thus hypothesized that NF10 mAb could affect SlpA renewal in the S-layer, thereby promoting bacterial lysis. To quantify cell lysis during exponential growth phase in the presence of the NF10 mAb, we monitored the lactate dehydrogenase (LDH) released, a strictly cytoplasmic enzyme used to analysis cell viability^26^. We found significantly more LDH in supernatants of NF10-exposed bacterial cultures compared to isotype control-exposed bacterial cultures (Fig. 3c), supporting the hypothesis that NF10 mAb weakens the integrity of the bacterial membrane.

If the bacterial membrane integrity is compromised, it should become vulnerable to enzymes, in particular to lysozyme. *C. difficile* strains are indeed highly resistant to lysozyme, a protein produced by Paneth cells in the small intestine and ascending colon in humans, while SlpA mutants’ growth is highly affected in the presence of lysozyme^5^. Strikingly, high concentrations of NF10 (100 and 200 μg/mL) only partially inhibited growth of *C. difficile* in standard culture conditions but abrogated growth in the presence of lysozyme (Fig. 3d, Supplemental Fig. 1c). Moreover, low concentrations of NF10 (6.25 μg/ml to 25μg/mL that did not affect growth in standard culture conditions significantly inhibited growth in the presence of lysozyme. The secondary bile acid deoxycholate (DCA) plays a major role in CDI^27^ and can abrogate at high dose, the growth of *C. difficile* bacteria^28^. ^28^ (i.e., 25 µg/mL) ^26^, that affect in standard culture conditions ^28^The presence of mAb NF10 significantly inhibit growth of *C. difficile* with subinhibitory concentrations of DCA^26^, even at concentrations of mAb insufficient to inhibit growth in standard culture conditions (Fig. 3d, Supplemental Fig. 1d). Altogether, these results show that mAb NF10 can potentiate by a synergistic effect, the detrimental effect of lysozyme or bile acid on *C. difficile* growth with.

### *C. difficile* toxin secretion is altered by anti-LMW mAbs

Even though *C. difficile* toxins are secreted through pores in the S-layer by a mechanism still incompletely known^4^, impaired toxin production has been reported in *C. difficile* SlpA-null mutants^5^. Consequently, we explored whether anti-LMW mAbs were able to alter toxin secretion *in vitro*. In our assay, CD630Δ*erm* secreted ∼18ng/mL at 24h and ∼170ng/mL at 48h of TcdA, and ∼1ng/mL at 24h and ∼14ng/mL at 48h of TcdB (Fig. 4). As expected, the Pathogenicity locus (Paloc)-deficient *C. difficile* mutant (ΔPaloc)^29^ that lacks the toxin A and toxin B genes did not secrete any detectable quantity of these two toxins. Incubation with mAb NF10, but not any other anti-LMW mAb, significantly increased TcdA and TcdB secretion at both 24h and 48h. In contrast, mAbs KH2 and TG10 significantly reduced TcdA and TcdB secretion at 48h. Surprisingly, mAb 2B7 that partially the same epitope as mAb KH2 (Fig. 1D) and has a better affinity for LMW (Table 1) did not affect the secretion of either toxin. Together, these results indicate that even though anti-LMW mAbs NF10, KH2 and TG10 bind the same target on the *C. difficile* surface, they induce contrasting effects on toxin secretion that appears tightly epitope-dependent.

**Figure 4:**
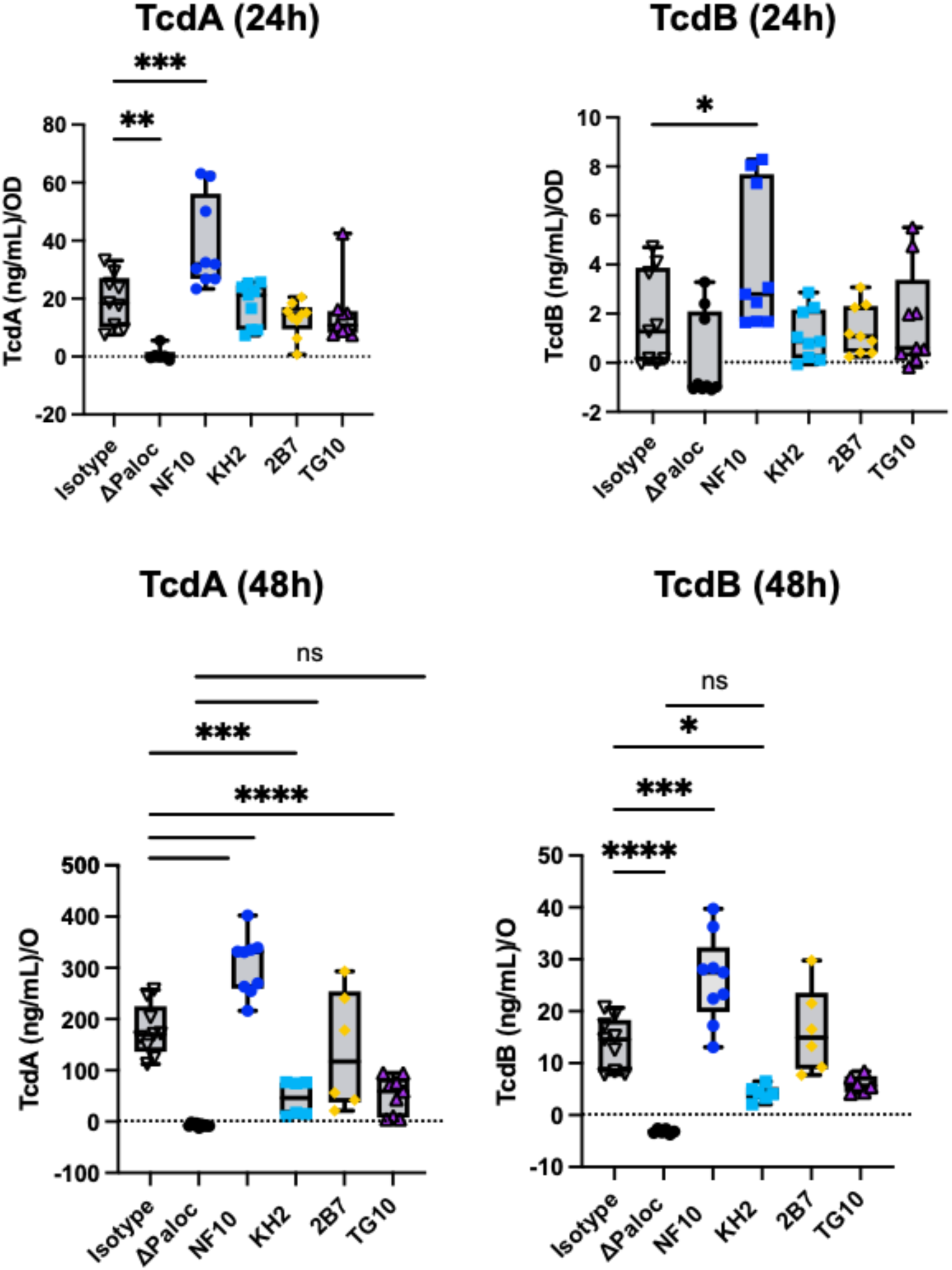
Anti-LMW mAbs modulate *C. difficile* toxin secretion. Quantification of TcdA or TcdB toxin secretion in CD630Δerm in the presence of anti-LMW mAbs or isotype control. CD630ΔermΔPaloc mutant strain has been tested as a negative control. Toxin titers in culture supernatants were determined at 24h and 48h by ELISA. Boxplots show medians (middle line) and interquartile range, and the whiskers indicate minimal and maximal values of three replicates. Asterisks indicate statistical significance calculated with a one-way ANOVA test followed by a Dunnett’s multiple comparison test (ns: not significant; * p < 0.05, * p< 0.01, *** p < 0.001, and **** p < 0.0001).

### *C. difficile* biofilm formation is increased by anti-LMW mAbs NF10 and 2B7

*C. difficile* CWP84 mutants with altered S-layer were reported to increase biomass of their biofilm compared to the parental strain, suggesting a role of SlpA in *C. difficile* biofilm formation^30^. We therefore assumed that biofilm formation could be modulated when *C. difficile* S-layer is constrained by anti-LMW mAbs. In presence of sub-inhibitory of DCA, the CD630Δ*erm* strain forms biofilm *in* ^28^and we showed that biomass of DOC-induced biofilm increases after mAb NF10 and mAb 2B7 incubation (Fig. 5a). By quantifying nucleic acids and proteins, we found that biofilm significantly increase after incubation with mAb 2B7, while a non-significant trend is observed after incubation with mAb NF10 (increase in biofilm formation compared to the wild type (100%): 175%, p=0.0231 and 149%, p=0.1661, for 2B7 and NF10 respectively; Fig. 5b). To strengthen these results, we analyzed biofilm volume, thickness and roughness (aka unevenness of the biofilm surface) using confocal laser scanning microscopy on fluorescently-labeled *C. difficile* as previously reported^31^. Incubation with either mAb NF10 or mAb 2B7 induced a ∼1.7-fold increase in biovolume, a ∼2-fold increase in thickness and a ∼1.6-fold increase in roughness when compared to DOC-induced biofilm in absence of mAbs (Fig. 5c-d). These results highlight the contribution of SlpA LMW in the biofilms formation, with epitope-dependent enhancement of biofilm generation revealed by two anti-LMW mAbs.

**Figure 5:**
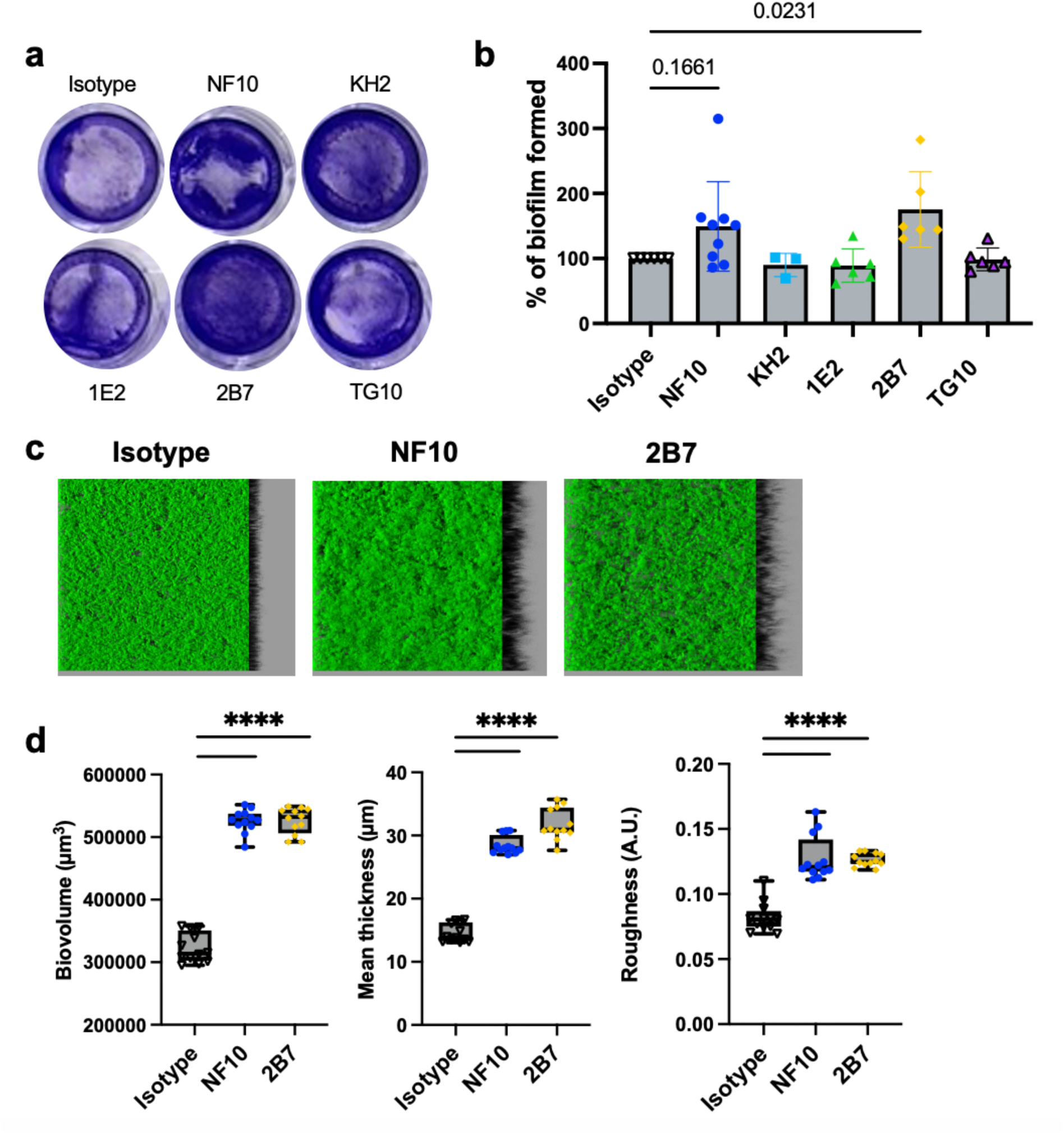
Anti-LMW mAbs influence *C. difficile* biofilm formation. Biofilm formation with CD630Δ*erm* strain was assayed in BHISG medium supplemented with 240 µM DCA. **a**. Representative pictures of biofilm formed in the presence of indicated mAbs after crystal violet staining. **b**. Biofilm biomass was assessed by absorbance at 600nm. Histograms show medians (middle line) and whiskers indicate standard deviation of at least three independent experiments. **c.** Visualization mAbs-coated CD630Δ*erm* biofilms stained with SYTO9. Z-stacks were analyzed with BiofilmQ. CLSM images are representative of three independent biological replicates. For each image, the virtual shadow projection of the biofilm is shown in dark on the right. **d**. Quantitative analyses were performed with BiofilmQ to measure the biovolume, thickness and roughness of the biofilms. The interquartile boxplots show medians (middle line) and the whiskers indicate minimal and maximal values of three replicative samples. Asterisks indicate statistical significance with a one-way ANOVA test followed by a Dunnett’s multiple comparison test (****p < 0.0001).

## Discussion

*C. difficile* is a complex pathogen to study, being anaerobic and lacking tools including antibodies, to investigate the contribution of its surface components to growth, adhesion, toxin secretion, infectivity, and biofilm generation among other of its properties. Herein, we identified the first series of anti-SlpA LMW mAbs and exploited them to demonstrate the contribution of LMW to growth, toxin secretion and biofilm formation, and its potential as a target for neutrophil-dependent phagocytosis. Interestingly, anti-LMW mAbs demonstrated various effects on *C. difficile* -sometimes opposite-depending on their epitope. Among them, the high-affinity anti-LMW mAb NF10 had multiple effects on *C. difficile* by impairing growth in a dose-dependent manner, increasing susceptibility to lysis by lysozyme and bile acid and affecting toxin secretion and biofilm formation. No such impact has been observed with the anti-LMW mAbs KH2 and TG10. However, contrary to the anti-LMW mAb NF10, they inhibit toxin secretion suggesting an epitope-dependent regulation of *C. difficile* biology by the low-molecular weight subunit of SlpA

One of the most surprising features of these anti-LMW mAbs is their contrasting effects depending on the epitope they bind to. Antibodies and nanobodies targeting *C. difficile* S-layer have been proposed as attractive therapeutic agents^16,32^. Likewise, active and passive immunization strategies have been tested with varying degrees of success to prevent or treat CDI^15,33^. Our findings suggest that anti-S-layer polyclonal responses include both beneficial and detrimental antibodies. Thus, the precise definition of the epitopes recognized of the S-layer associated to their effect on various *C. difficile* functions is of the outmost importance to design successful anti-S-layer therapeutic agents. Furthermore, even if a toxin-suppressing antibody might at first glance appear beneficial to the host, it might also facilitate biofilm formation and therefore can promote gut persistence of *C. difficile*. Our data prompt to test novel therapeutic agents not only on single episode CDI models, but also on recurrence models, to evaluate the role of the biofilms as a reservoir for further infections.

The S-layer is an important component involved during bacterial growth since new S-layer must be continuously assembled when cells divide. While no previous study could evaluate the effect of targeting the *C. difficile* S-layer due to the lack of specific antibodies, a related study on *Bacillus anthracis* showed that anti-S-layer nanobodies attenuated bacterial growth^34^, reminiscent of our findings with mAb NF10 on the growth of *C. difficile*. The authors showed that nanobodies inhibited S-layer *de novo* assembly with a full dissolution of S-layer polymers, which resulted in drastic morphological defects and S-layer disruption. In the same way, mAb NF10 may also prevent optimal S-layer compaction leading to morphological defects and bacterial lysis. On the contrary, *C. difficile* S-layer null mutants did not show any growth defects^5^, but were more susceptible to lysozyme and anti-microbial peptides such as LL-37. Consistently, we showed in our work that addition of mAb NF10 leads to *C. difficile* sensitivity to lysozyme. As shown by Salgado *et al., C. difficile* S-layer forms a tightly compact barrier around the bacteria, impenetrable to large molecules^4^. Besides S-layer disruption, we propose that mAb NF10 interaction with *C. difficile* LMW could enable the import of large molecules *e.g*., lysozyme (14kD), a promising process which could be used for specific drug delivery.

Toxin secretion is a major physiological process that confers its pathogenicity to the bacteria. Since CDI symptoms are mainly due to TcdA and TcdB production, the regulation of their expression and the mechanisms involved in their secretion have been extensively studied^35^. Toxin synthesis is growth phase-dependent and regulated in response to a variety of environmental factors such as availability of specific nutrients, temperature, and cell density^35–38^. While toxin secretion is depending of an holin-depending system,^39^ how the toxins cross the *C. difficile* membrane and consequently how they interact with the S-layer without bacterial lysis remain open questions^4^. S-layer must create discrete pores to allow toxin export while maintaining bacterial integrity. Interestingly, three of our anti-LMW mAbs alter toxin secretion: one by increasing it, the other while two by decreasing it, pointing towards a dual role of S-layer in toxin release. On the one hand, S-layer disruption by mAb NF10 may lead to a massive toxin release, on the other hand mAbs KH2 and TG10 may “rigidify” or “lock” the S-layer, thus abrogating toxin export. Consistent with our findings, mutants affecting *C. difficile* S-layer displayed these contrasting features^5,40,41^. We may also hypothesize that changes in the S-layer integrity may modulate toxin expression. Further functional and structural studies are needed to solve how SlpA impacts on import-export mechanisms *in C. difficile*.

Another aspect of *C. difficile* pathogenicity relies on its ability to forms biofilms, which been suggested to contribute to the pathogenesis and persistence of *C. difficile*^42^. Indeed, biofilm-like structures have been observed in CDI mouse models *in vivo*^43,44^. Analyses of *C. difficile* biofilm composition showed that extracellular DNA is an essential component of the biofilms matrix. Of note, incubation with DNase I drastically reduced the biofilm biomass^45,46^. These data are in agreement with our hypothesis that mAb NF10-induced lysis facilitates biofilm formation by increasing the amount of extracellular DNA and proteins in the biofilm matrix. Beyond S-layer disruption and bacterial lysis, the extent to which S-layer proteins such as LMW are *per se* involved in biofilm formation remains unclear. Inhibition of S-layer-mediated aggregation could also impact the early steps of biofilm formation, as has been demonstrated for *Lactobacillus helveticus M92*^47^.

Our study has limitations. We studied biofilm formation and architecture in a closed system with one *C. difficile* strain. As a recent study demonstrated that biofilms grown in well-plates and biofilms obtained in open systems harbor different characteristics in terms of cell-surface protein expression^48^, it would be judicious to evaluate anti-LMW mAbs in other biofilm forming conditions. Moreover, biofilm-forming ability differs between *C. difficile* strains^49^, making difficult to assign our model to all ribotypes. Besides, knowing the precise LMW epitopes that are recognized by the mAb series we describe here could help to decipher the varying effects these have on *C. difficile* physiology. Secretory IgA have indeed been reported to shape functional microbial fitness depending on the recognized antigen and epitopes^50^. The absence of the D2 domain of the LMW in *C. difficile* has been shown to be sufficient to confer susceptibility to lysozyme, therefore indicating its crucial role in maintaining S-layer integrity^4^. We hypothesize that mAb NF10 interacts with an epitope in the D2 domain, thus impairing its function and therefore mimicking what has been found with the mutant lacking this domain.

In this work, we demonstrate that targeting of mAbs to the S-layer of *C. difficile* has multiple and contrasting effects on the physiology of the bacteria. This study provides insights on the function of the *C. difficile* S-layer and suggests ways to target and modify some of its physiological processes. Future fine-tuned work on mAbs recognizing a determined epitope on the S-layer, leading to a precise function such as impaired growth or decrease in toxin secretion, could lead to new therapeutic strategies for CDI.

## Methods

### Production of recombinant LMW proteins

Recombinant *C. difficile* LMW-630 was produced as C-terminal 6xHis-tagged proteins from plasmid pET-28a(+) (TwistBiosciences, #69864). Plasmids were transformed into *Escherichia coli* strain D43 and grown in NZY auto-induction lysogeny broth (LB) medium (NZYtech, #MB180). Bacteria were harvested by centrifugation and lysed using Precellys system according to manufacturer instructions (Bertin Technologies, #P002511-PEVT0-A.0). Recombinant LMW-SLP proteins from the soluble fraction were purified by affinity chromatography on Ni-agarose columns using an AKTA prime (GE Healthcare, #11001313). Proteins were dialyzed against 10 mM HEPES pH 7.5, 150 mM NaCl prior to analysis or long-term storage.

### Generation of monoclonal antibodies against LMW of *C. difficile* strain 630

Knock-in mice expressing human antibody variable genes for the heavy (VH) and kappa light chain (Vκ) (VelocImmune) were described previously^17,18^ and provided by Regeneron Pharmaceuticals to be bred at Institut Pasteur. BALB/c mice were purchased from Janvier labs. All animal care and experimentation were conducted in compliance with the guidelines. The study, registered under #210111 was approved by the Animal Ethics committee CETEA (Institut Pasteur, Paris, France) and by the French Ministry of Research.

BALB/c mice and VelocImmune mice were injected at day 0, 21 and 42 with 50 μg of recombinant LMW630 mixed with 200 ng/mouse pertussis toxin (Sigma-Aldrich, MO, USA). Enzyme-linked immunosorbent assay was performed to measure serum responses to antigen (see methods below) and the 3 best immunized animals were boosted with the same mix. Four days later, splenocytes were fused with myeloma cells P3X63Ag8 (ATCC, France) using ClonaCell-HY Hybridoma Kit according to manufacturer’s instructions (StemCell Technologies, Canada). Culture supernatants were screened using ELISA (see below) and antigen-reactive clones were expanded in serum IgG free RPMI-1640 (Sigma-Aldrich, MO, USA) into roller bottles (Sigma-Aldrich, MO, USA) at 37°C. After 14 days, supernatants were harvested by centrifugation at 2500 rpm for 30 min and filtered (0.2 µm). Antibodies were purified by protein A affinity chromatography (AKTA, Cytiva, Germany) as described previously^51^.

### ELISA assays

Maxisorp microtiter plates (Dutscher, France) were coated with 0.3 μg of LMW630 recombinant protein in carbonate buffer (Na_2_CO_3_/NaHCO_3_) for 2 hours at room temperature (RT). Free sites were blocked by a 2-hour incubation at RT with 1X-PBS 1% BSA. Plates were washed three times with 1X-PBS 0.05% Tween 20 (PBS-T) before being co-incubated with serum, supernatants or mAbs at different concentrations (from 10^-6^ μg/mL to 10μg/mL) for 1h at RT. After five washes, goat anti-mouse IgG Heavy and Light Chain antibody HRP-conjugated (Bethyl, TX, USA, dilution 1:20 000) was added for 1h at RT followed by incubation with OPD substrate revealing reaction for 10 min (Sigma-Aldrich, MO, USA). Absorbances were analyzed at 495 *vs* 620 nm on an ELISA plate reader (Berthold, France).

### Bio-layer interferometry

Biolayer interferometry assays were performed using Anti-Mouse IgG Fc Capture biosensors (18-5088) in an Octet Red384 instrument (ForteBio, USA). MAbs (10 μg/mL) were captured on the sensors at 25°C for 1800 seconds. Biosensors were equilibrated for 10 minutes in 1X-PBS, 0,05% Tween 20, 0.1% BSA (PBS-BT) prior to measurement. Association was monitored for 1200s in PBS-BT with LMW630 at a range of concentrations from 0.01 nM to 500 nM followed by dissociation for 1200s in PBS-BT. For epitope competition assays, sensors were further immersed in solutions containing mAbs at 10 μg/mL. Biosensor regeneration was performed by alternating 30s cycles of regeneration buffer (glycine HCl, 10 mM, pH 2.0) and 30s of PBS-BT for 3 cycles. Traces were reference sensor (sensors loaded with an irrelevant mAb) subtracted and curve fitting was performed using a global1:1 binding model in the HT Data analysis software 11.1 (ForteBio, USA), allowing to determine K_D_ values.

### IgH and IgL sequencing

Total RNA was extracted from murine splenocytes using NuceloSpin RNA plus kit (Macherey-Nagel, France) according to the manufacturer’s instruction. cDNA were generated at 50°C for 60 min using random primers and SuperScript III Reverse Transcriptase (Invitrogen, MA, USA). The primer pairs for IgH and IgL, described in Supplemental Table 2 were used for amplification with GoTaq G2 polymerase (Promega, WI, USA). Amplification was performed by 35 cycles PCR each consisting of 94°C for 30 sec, 63°C for 30 sec, 72°C for 30 sec. At the end of the 35 cycles, samples were run for an additional 10 min at 72°C and analyzed by 1.5% agarose gel electrophoresis. PCR products were then sequenced by Eurofins (France) using 3’ primers.

### Flow cytometry assay

mAb binding to whole bacteria was assessed by bacterial flow cytometry assays, as previously described^20^. Briefly, fixed *C. difficile* (10^6^/condition) were stained with 5 μM SYTO9 dye (Thermo Fisher Scientific, MA, USA) in 0.9% NaCl for 30 min at RT. Bacteria were washed (10 min, 4000g, 4°C) and resuspended in 1X PBS, 2% BSA and 0.02% Sodium Azide (PBA). Mabs were pre-diluted in PBA at 20 µg/mL and incubated for 30 min at 4^◦^C. Bacteria were washed, and AF647 AffiniPure goat anti-mouse IgG (H+L) antibody or isotype control (dilution 1:200, Jackson ImmunoResearch, PA, USA) were incubated for 30 min at 4^◦^C. After washing, bacteria were resuspended in sterile 1X-PBS. Flow cytometry acquisition was performed on a MacsQuant cytometer (Miltenyi, Germany) and analyzed on FlowJo software (BD Biosciences, CA, USA).

### Isolation of human neutrophils

Human peripheral blood was collected on EDTA from healthy volunteers. Blood neutrophils were separated by negative magnetic selection (MACSxpress, Miltenyi Biotec, Germany) according to the manufacturer’s instructions. After negative selection, the neutrophil-enriched suspension was recovered, and residual erythrocytes were further removed using the MACSxpress Erythrocyte Depletion kit (Miltenyi Biotec, Germany). The resulting neutrophil suspension was washed with HBSS (Sigma-Aldrich, MO, USA) and resuspended to an appropriate volume in HBSS (Ca^2+^/ Mg^2+^) + 2% fetal calf serum (Cytiva, Germany).

### Phagocytosis assay

Human neutrophils were plated at a concentration of 8 x 10^5^ cells/ml. Fixed *C. difficile* were incubated with one mAb at 20 µg/mL or a cocktail of mAbs NF10, KH2, 1E2, 2B7 and TG10 at equimolar ratio and stained with pHRodo dye (Thermo Fisher Scientific, MA, USA) following the manufacturer instructions. Mouse anti-rocuronium mAb (in house production) was used as isotype control. Bacteria were then incubated with neutrophils at a Multiplicity Of Infection (MOI) of 100 for 1.5h at 37°C (20,000 neutrophils for each condition). Flow cytometry acquisition was performed on a MacsQuant16 cytometer (Miltenyi, Germany) and analyzed on FlowJo software v10.8.1 (BD Biosciences, CA, USA).

### Bacterial strains and culture conditions

*C. difficile* 630Δerm^52^, a spontaneous erythromycin sensitive derivative of the reference strain 630, and *C. difficile* strain UK1^53^ of ribotype 027 strains were grown anaerobically (5% H2, 5% CO2, 90% N2) in TY medium (30 g/L tryptone, 20 g/L yeast extract) or in Brain Heart Infusion (BHI) medium supplemented with 0.5% (w/v) yeast extract, 0.01 mg/mL cysteine and 100 mM glucose (BHISG). All media and chemicals were purchased from Sigma-Aldrich, MO, USA.

### Growth assays, lysozyme resistance and quantification of lysis

Overnight *C. difficile* cultures were grown in TY broth, subcultured to an Optical Density at 600 nm (OD600nm) of 0.05 in 200 µL of BHISG or, when appropriate, BHISG supplemented with DCA (240 µM, Sigma-Aldrich, MO, USA) in 96-well flat bottom plates (Merck, Germany) and then grown for 24h or 18h with OD600nm measurements every 30 min taken by GloMax Plate Reader (Promega, WI, USA). Anaerobiosis was maintained with a O_2_-less sealing film (Sigma-Aldrich, MO, USA). Where appropriate, lysozyme (1 mg/mL) was added after 2.5h of growth. Experiments were performed at least in triplicate. For lysis quantification, LDH was measured in 13h-culture supernatants using CytoTox 96 Non-Radioactive cytotoxicity assay according to manufacturer instructions (Promega, WI, USA).

### Biofilm assays

Overnight cultures of *C. difficile* 630Δerm grown in TY medium were diluted to 1:100 into fresh BHISG supplemented or not with 240 µM DCA and 0.2 mg/mL mAbs. 1 mL of diluted cultures were added in 24-well plates (polystyrene tissue culture-treated plates, Costar, USA). Then, plates were incubated at 37°C in an anaerobic environment for 48h. Biofilm biomass was measured using an established method^28^. Briefly, biofilms were washed with 1X-PBS and stained with crystal violet for 5 min. After washing, crystal violet was resuspended in ethanol and OD_600nm_ measured.

### Confocal Laser Scanning Microscopy (CLSM)

Biofilms were grown in 96-well plates (Microclear, Greiner Bio-one, France) in BHISG supplemented with DCA (240 μM) and anti-LMW630 mAbs as described above. After 48h, supernatants were carefully removed by pipetting and biofilms were fixed with 4% paraformaldehyde (Sigma-Aldrich, MO, USA). Biomass was then stained with SYTO9 dye (Life Technologies, USA) at a final concentration of 20 μM. Dye were incubated for 30 min before CLSM imaging/analysis. Z-stacks of horizontal plane images were acquired in 1 μm steps using a Leica SP8 AOBS inverted laser scanning microscope (CLSM, LEICA Microsystems, Wetzlar, Germany) at the INRAE MIMA2 platform (doi.org/10.15454/1.5572348210007727E12)^54^.^54^. At least two stacks of images were acquired randomly on three independent samples at 800 Hz with a x63 water objective (N.A.=1.2). Fluorophores were excited, then their emissions were captured as prescribed by the manufacturer.

### Analysis of CLSM biofilm images

Z-stacks from the CLSM experiments were analyzed with the BiofilmQ software^55^ to extract quantitative geometric descriptors of biofilms structures. Images were all treated with the same process in each fluorescence channel. First, the images were denoised by convolution (dxy=5 and dz=3), then they were segmented into two classes with an OTSU thresholding method with a sensitivity of 2. The detected signal was then declumped in 3.68 μm cubes and small objects were removed with a threshold of (0.5μm^3^) to clean the remaining noise. Exported data were analyzed in the software Imaris to generate biofilm 3D projections and in GraphPad prism to generate quantitative graphs.

### Toxin A & B assays

*C. difficile* 630Δerm and 630ΔermΔPaloc were grown in 6-well plates containing 2 mL of TY medium for either 24h or 48h. Absorbances at 600 nm were measured, then cultures were harvested and centrifuged at 4,000 rpm for 5 min. Toxins were assessed in supernatants using ELISA. Maxisorb microtiter plates (Dutscher, France) were coated with 5 μg/mL of anti-TcdB capture antibody (BBI solutions, Madison, WI) or anti-TcdA capture antibody (Novus Biological, CO, USA). Purified toxin A and B (Sigma-Aldrich) were used as standards. Supernatants were added for 1h30 at RT. After washing, anti-toxin B biotinylated antibody (BBI solutions, Madison, WI) followed by high sensitivity Streptavidin-HRP conjugate (ThermoFisher, Waltham, MA), or anti-toxin A HRP-conjugated antibody (LSBio, WA, USA) signal was detected with TMB substrate (ThermoFisher, Waltham, MA) at 450nm using a ELISA plate reader (Berthold, France). Toxin concentrations were normalized with OD_600nm_ values for each well.

### Statistical analysis

Growth, LDH, toxins and biofilm’ assays values were analyzed in Prism 8.0 (GraphPad, San Diego, CA). Statistical analysis was performed using one-way ANOVA test followed by a Dunnett’s multiple comparison test. A p value ≤0.05 was considered significant.

## Acknowledgements

This work was funded by Fondation Janssen Horizon, the Institut National de la Santé et de la Recherche Médicale (INSERM) and the Institut Pasteur. LH is a doctoral fellow of Sorbonne Université. DS was partly supported by a poste d’accueil 2017 Institut Pasteur – Assistance Publique des Hôpitaux de Paris (APHP) and by the Agence Nationale de la Recherche (ANR) program Résilience-Covid-19 MUCOVID. Work in the G. Gorochov’s team is supported by Institut National de la Santé et de la Recherche Médicale (INSERM), Sorbonne Université, Fondation pour la Recherche Médicale (FRM), Paris, France, program ‘‘Investissement d’Avenir’’ launched by the French Government and implemented by the Agence Nationale de la Recherche (ANR) with the reference COFIFERON ANR-21-RHUS-08, by EU Horizon HLTH-2021-DISEASE-04 UNDINE project, by programme DIM Ile de France thérapie cellulaire et génique, by Fondation pour la Recherche Médicale, Paris, France (Programme Equipe FRM 2022) and by the Département Médico-Universitaire de Biologie et Génomique Médicales (DMU BioGen), APHP, Paris, France.

---

**Supplemental Figure 1.**
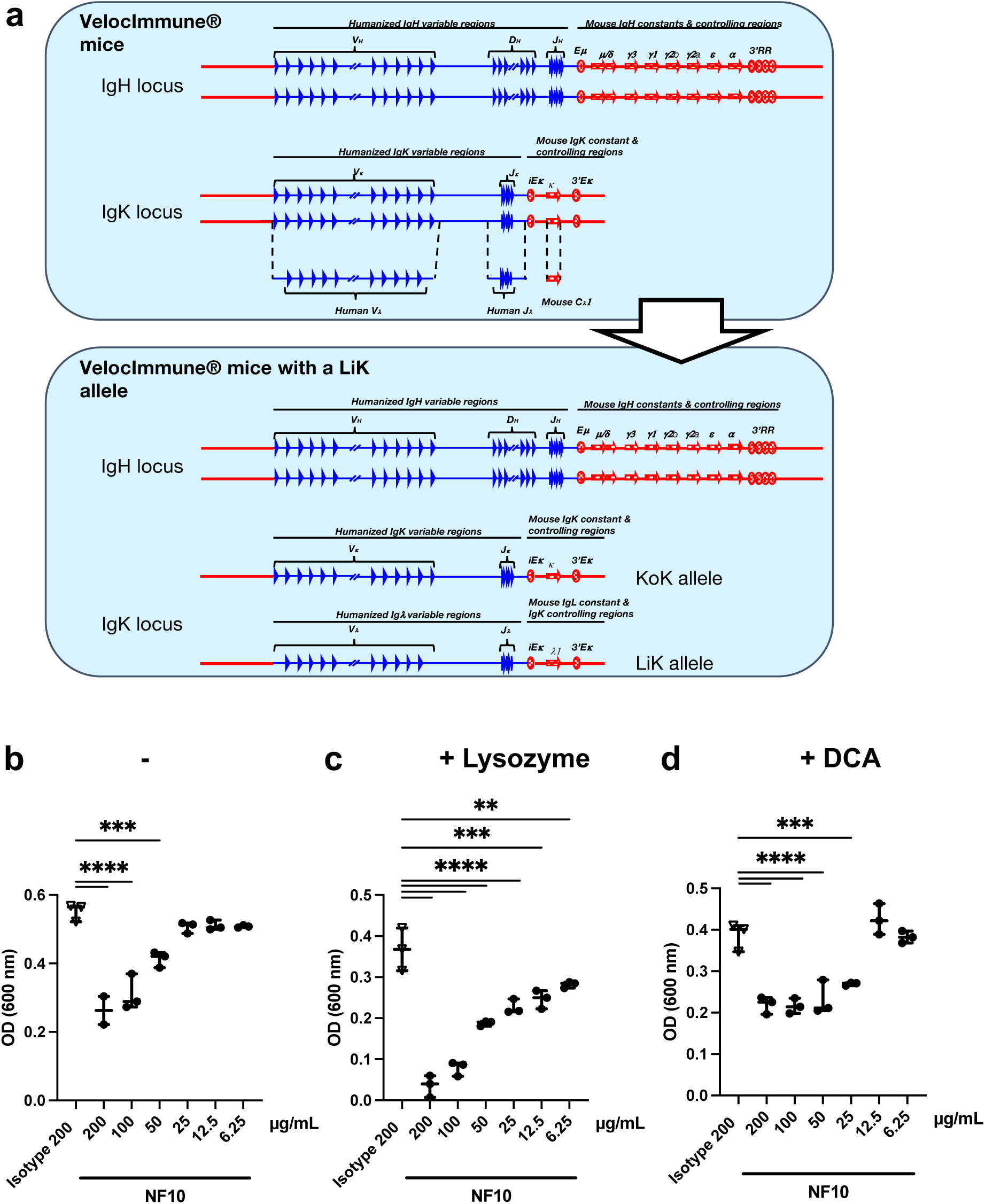
**(a)** Schematic of the generation of mice knock-in for the human variable VDJ segments in the endogenous variable heavy chain locus, and for the human variable VJ segments in the endogenous variable light chain kappa locus. **(b-d)** Cultures of *C. difficile* 630Δerm incubated with different concentrations of NF10 mAb were monitored in the absence (**b**) or in combination with either (**c**) lysozyme (500 μg/ml), which was added after 2.5h growth or (**d**) DCA (240 µM). Isotype control was included in all experiments. (**b-d**) The boxplots show medians (middle line) and the whiskers indicate min and maximal values at 13 hours. Asterisks indicate statistical significance with a one-way ANOVA test followed by a Dunnett’s multiple comparison test (ns: not significant; * p < 0.05, ** p < 0.01, *** p < 0.001, and **** p < 0.0001).

**Supplemental Table 1:** Ig gene analysis and kinetic parameters of anti-LMW mAbs. V(D)J families were obtained by blasting the sequences on IMGT data base and kinetic parameters determined using the BLI analysis software. ND: not determined.

| mAbs | Affinity |  |  |
| --- | --- | --- | --- |
|  | Kd (pM) | kon (1/Ms) | koff (1/s) |
| NF10 | 43 | 7,00E+05 | 3,00E-05 |
| KH2 | 172 | 1,31E+05 | 2,25E-05 |
| 1E2 | 176 | 2,00E+05 | 3,53E-05 |
| 2B7 | 67 | 1,42E+06 | 9,46E-05 |
| 2C4 | 1370 | 4,48E+05 | 3,23E-04 |
| 4G4 | 70000 | 2,22E+05 | 1,56E-02 |
| TG10 | 4200 | 3,02E+05 | 1,22E-03 |

**Supplemental Table 2:**
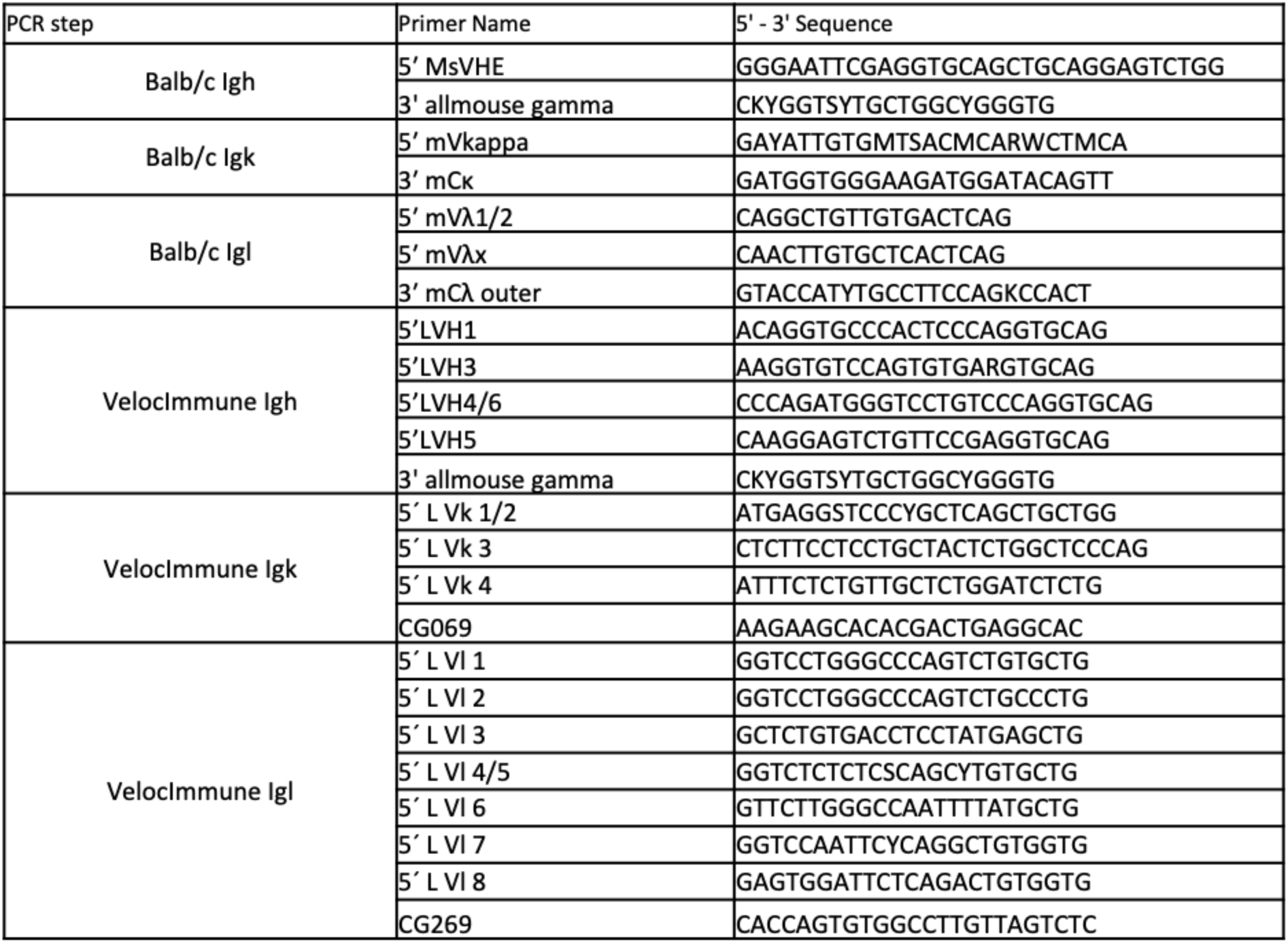
Primers for Ig gene amplification of BALB/c and VelocImmune mice.

